# Transcriptional network of the industrial hybrid *Saccharomyces pastorianus* reveals temperature-dependent allele expression bias and preferential orthologous protein assemblies

**DOI:** 10.1101/2021.04.07.438844

**Authors:** Soukaina Timouma, Laura Natalia Balarezo-Cisneros, Javier Pinto, Roberto De La Cerda, Ursula Bond, Jean-Marc Schwartz, Daniela Delneri

## Abstract

*Saccharomyces pastorianus* is an industrial natural yeast evolved from different hybridisation events between the mesophilic *S. cerevisiae* and the cold-tolerant *S. eubayanus*. This complex aneuploid hybrid carries multiple copies of the parental alleles alongside specific hybrid genes and encodes for multiple protein isoforms which impart novel phenotypes, such as the strong ability to ferment at low temperature. These characteristics lead to agonistic or antagonistic competition for substrates and a plethora of biochemical activities, resulting in a unique cellular metabolism. Here, we investigated the transcriptional signature of the different orthologous alleles in *S. pastorianus* during temperature shifts. We identified temperature-dependent media-independent genes and showed that 35% have their regulation dependent on extracellular leucine uptake, suggesting an interplay between leucine metabolism and temperature response. The analysis of the expression of ortholog parental alleles unveiled that the majority of the genes express preferentially one parental allele over the other, and that *S. eubayanus*-like alleles are significantly over-represented among the genes involved in cold acclimatisation. The presence of functionally redundant parental alleles may impact on the nature of protein complexes established in the hybrid, where both parental alleles are competing. Our expression data indicate that the majority of the protein complexes in the hybrid are likely to be either exclusively chimeric or uni-specific, and that the redundancy is discouraged, a scenario which fits well with the stoichiometric balance-hypothesis. This study offers the first overview of the transcriptional pattern of *S. pastorianus* and provide a rationalisation for its unique industrial traits at expression level.

## Introduction

*Saccharomyces pastorianus* is a natural evolved allopolyploid and sterile hybrid between the mesophilic *Saccharomyces cerevisiae* and the cold-tolerant *Saccharomyces eubayanus. S. pastorianus* is used in 89% of brewed beer worldwide (Gorter de Vries *et al*. 2019). This species has been isolated, domesticated and maintained by human selection for colder brewing temperatures of 8-15°C (lager fermentation environment) (Nakao *et al*. 2009). Before the discovery of *S. eubayanus* species, the non-*S. cerevisiae* portion of the genome of *S. pastorianus* was considered as being *S. uvarum* and/or *S. bayanus* genome, that are closely related to *S. eubayanus* (Vaughan-Martini and Martini 1993). *S. cerevisiae* was isolated in Europe, while *S. eubayanus* have been isolated from Nothofagus trees in Patagonia and in East Asia (Tibet) (Bing *et al*. 2014). The connection of Asia and Europe via the silk road could explain how the hybridisation occurred between those two species. Strong evidence suggests that the ancestry of *Saccharomyces sensu stricto* yeast underwent an event of whole genome duplication (WGD), that may have occurred 100 million years ago (Wolfe 2015). The doubling of an organism’s genetic content leads immediately to a reproductive barrier with their relative species and ancestors. However, in the yeasts, two ancestral species mated before the WGD which restored its fertility. WGD is a rare evolutionary event with great consequences as new paralogs are the basis of speciation, as they can sub-functionalise. The analysis of transposon sequence distribution within sub-telomeric regions and recombination breakpoints in the genome of *S. pastorianus* strains suggested its separation into two genomically distinct groups, which may have arisen from multiple hybridisation events (Monerawela and Bond 2018). One hypothesis state that an initial spontaneous hybridisation event occurred between a haploid *S. cerevisiae* strain and a diploid *S. eubayanus* strain (Bond *et al*. 2004). That event led to a progenitor of the Group I strains, which evolved through further reduction of the *S. cerevisiae* genome content, to produce an aneuploid and an approximate triploid genome. In parallel, the progenitor strain made a second hybridisation event with a different *S. cerevisiae* strain and evolved to give strains with an approximate tetraploid genome, that are classified in the Group II. A more recent theory supports that a single hybridisation event occurred between a diploid *S. cerevisiae* and a diploid *S. eubayanus*, resulting in a tetraploid progenitor that evolved to give the Group II strains (Okuno *et al*. 2016). In parallel, this progenitor underwent chromosomal deletions of the *S. cerevisiae* sub-genome, leading to a progenitor approximately triploid, that evolved to give the Group I strains (Salazar *et al*. 2019, Alsammar and Delneri 2020). Therefore, Group II strains possess about 2 to 3 times more *S. cerevisiae* genomic content than the Group I strains.

The difference in the genome composition of Group I and Group II strains results in different brewing properties. Group I strains appears to be more cold-tolerant (*i*.*e*., *S. eubayanus* trait) while Group II strains show a better maltotriose consumption (*i*.*e*., *S. cerevisiae* trait) (Brouwers *et al*. 2019). Group I include Saaz-type and Carlsberg type strains from Czech Republic and Denmark breweries, respectively. Group II, also referred to Frohberg-type, includes strains brewed in Canada, the Netherlands (Heineken, Oranjeboom and other breweries), and Denmark (non-Carlsberg breweries) (Dunn and Sherlock 2008). The *S. cerevisiae* parental strain of the Group I yeasts is related to those used for Ale beer production in Europe, while the one of the Group II contains, in addition to the Ale-like gene content, Stout-like sub-genome. *S. eubayanus* parental strain possesses similar fermentation capacities to Saaz group at cold temperature (10°C), and are more resistant to cold temperatures compared to Frohberg group and *S. cerevisiae* Ale strains (Gibson *et al*. 2013). *S. pastorianus* has also the ability to ferment under stressful conditions such as anaerobiosis, high hydrostatic pressure and high gravity sugar solutions, producing complex metabolites leading to unique flavours and aromas (Monerawela and Bond 2017a). The parental chromosomes of *S. pastorianus* underwent homeologous recombination, which resulted in unique hybrid chromosomes (Monerawela and Bond 2017b). Therefore, multiple copies of *S. cerevisiae*-like, *S. eubayanus*-like and hybrid gene alleles coexist in the genome. Consequently, the presence of multiple protein isoforms can lead to agonistic or antagonist competition for substrates. Protein redundancy can also broaden the possibilities of complex composition, that can be either uni-specific (*S. cerevisiae*-like or *S. eubayanus*-like) or potentially chimeric (Piatkowska *et. al* 2013). This mixture of proteomes results in flexible biochemical activities leading to a unique cellular metabolism. The *S. cerevisiae*-like and *S. eubayanus*-like alleles of three *S. pastorianus* Group I strains (CBS 1513, CBS 1503 and CBS 1538) and one Group II strain (WS34/70) have been predicted using HybridMine (Timouma *et al*. 2020), an open-source predictive tool our group developed (available on GitHub, https://github.com/Sookie-S/HybridMine).

Here, we focused on the Group I *S. pastorianus* CBS 1513, also known as *S. pastorianus carlsbergensis*, which is the first established lager yeast strain (Hewitt et al., 2014; Walther *et al*. 2014). This bottom fermenting strain has been used in breweries for lager-style production since its isolation by Emil Chr. Hansen in 1883 (Hansen, 1883). The brewing and distilling industries are constantly looking to improve their production strains for specific properties such as temperature tolerance, maltose utilisation, and balanced flavour profiles (Gorter de Vries *et al*. 2020). In *Saccharomyces*, a large number of studies presents temperature as a factor that influences several traits, including fermentation, cell division or protein synthesis (Hartwell and McLaughlin 1968, Ciani *et al*. 2016). At low temperature (<15°C), the beer fermentation process is slowed down, which causes a positive impact on volatile flavour production and retention, and increases the consumption of the wort sugars (García-Ríos *et al*. 2019). However, low temperatures can also be responsible for the termination of the fermentation process. Hybrids of cryo- and thermo-tolerant strains like *S. pastorianus* have a favourable genetic background to ferment at different temperatures. Historically, temperature treatments on *Saccharomyces* yeast had been used for industrial purposes, with beer brewing being one of the main applications. In this study, we analysed the transcriptome data of *S. pastorianus* CBS 1513 at different environmental conditions to determine the genes involved in temperature stress that are media-independent. We observed the presence of a skew towards the expression of one or the other parental sub-genome (*S. cerevisiae* or *S. eubayanus*) in the different conditions. Using the genomic features and expression data, we predicted the proportion of protein complexes that are more likely to be chimeric, uni-specific, fully or partially redundant in *S. pastorianus* CBS 1513. We identified that the majority of the protein complexes tend to be either chimeric or uni-specific even if both parental alleles are present in the genome. This study highlights that functional redundancy is efficiently exploited by *S. pastorianus* by using preferentially one allele over the other to adapt its environment. As this strain is still evolving, most of the redundant unused alleles could be predicted to be eventually lost. Our study is the first to provide insights into the genetics of *S. pastorianus* CBS 1513 at gene expression level and a rationalisation for its unique industrial traits.

## Results and Discussion

### 1. Temperature variations induce larger transcriptional changes than nutritional media

To determine the genes affected by temperature stress in different culture media, the expression profiles of the *S. pastorianus* CBS 1513 (Group I strain) at 13°C, 22°C and 30°C in SD media, wort (maltose rich medium), SD media with 6% ethanol, and SD media without leucine were analysed. These media were chosen given their relevance to industrial conditions, as wort is used during the beer fermentation, alcohol is produced during this process, and the leucine amino acid metabolism is responsible for important flavour compound production, such as the isoamyl-acetate production (fruity flavour) (Stewart 2017). The cells were grown aerobically and the yeast growth on these media and their comparison is reported in Figure S1, Tables S1 and S2. The expression of 6 *S. cerevisiae*-like and 6 *S. eubayanus*-like alleles was validated by RT-qPCR (Figure S2).

To find patterns of expression or groupings between the different conditions a multi-dimensional scaling (MDS) analysis of the RNAseq libraries was performed (Figure 1). When comparing the temperatures with the media, as expected, we observed that biological replicates clustered tightly together. The highest amount of variance is detected at different temperatures, where samples taken at the lowest temperature clusters at the opposite end to those at a highest temperature (*i*.*e*., larger spread of data on the first dimension). In a consistent trend, samples taken at 22°C clustered in between. The media show less variance with the exception of SD +6% ethanol at 30°C. The combined presence of ethanol and high temperature may increase the stress in the cell compared to the other conditions, which could explain the variance in the data. Overall, these results suggest that temperature stress has a higher effect on the transcriptome than the nutritional media and that temperature impacts on the growth of *S. pastorianus* in all media (Table S1).

**Figure 1:**
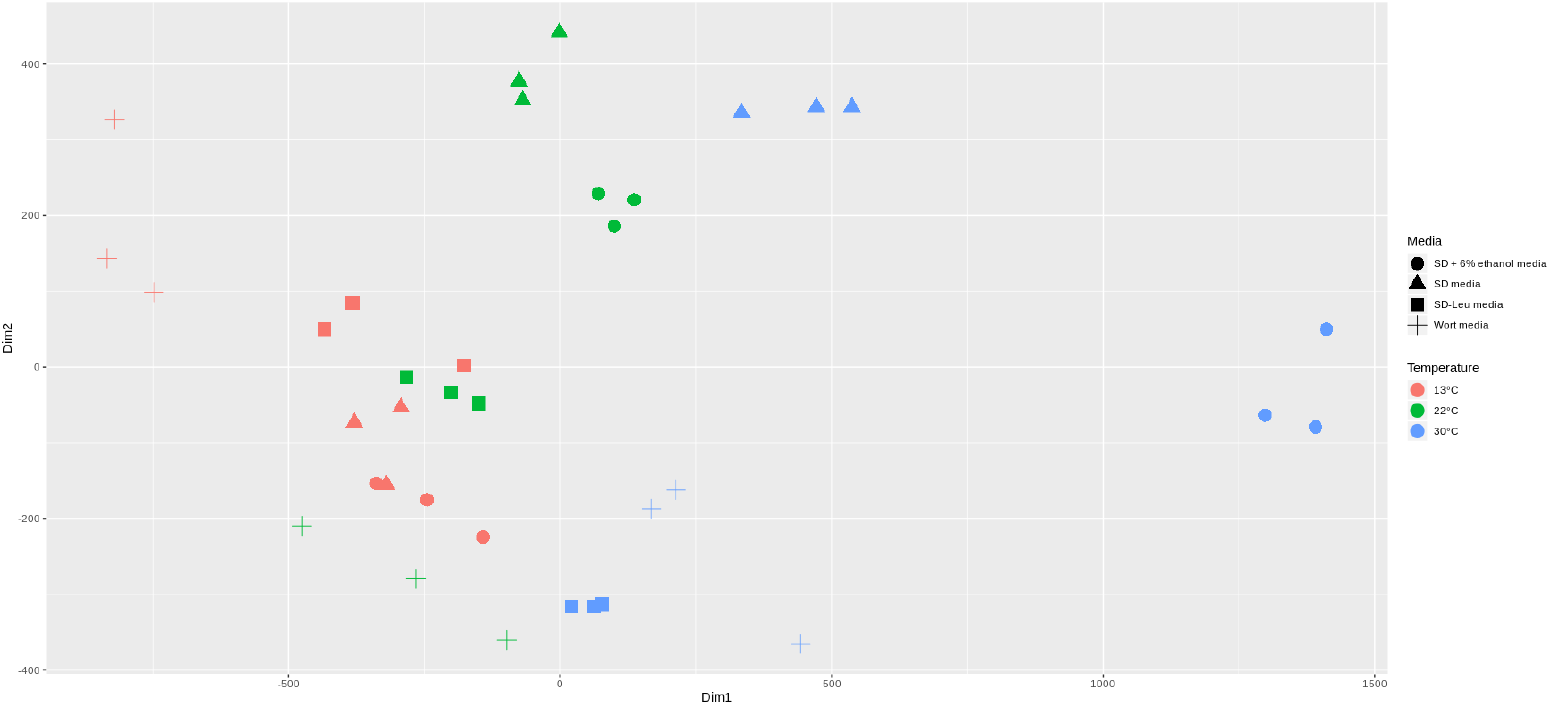
Multi-dimensional scaling (MDS) plot of RNA-seq expression profiles of *S. pastorianus* CBS 1513, in two dimensions. For visual aid, samples cultured at 13°C, 22°C and 30°C are coloured in red, green and blue, respectively. Samples cultured in the different media are represented with circles, triangles, squares and crosses shapes for SD + 6% ethanol, SD with all amino acids, SD w/o leucine and wort media, respectively.

To assess whether the type of culture media had an impact on the expression network at different temperatures, we compared the absolute number of differentially expressed (DE) genes (absolute fold change > 2 and p-adjusted value <0.05) obtained in each temperature condition, for each media. The absolute number of DE genes up- and down-regulated at different temperatures varied significantly according to the culture media (Figure 2). As expected, at extreme temperature conditions (13°C vs 30°C), changes in the transcriptome are higher in each culture media compared to the other temperatures (13°C vs 22°C and 22°C vs 30°C), with the exception of wort where the total number of DE genes is similar for all temperatures (Figure 2). Within the SD media, there are less DE genes at 22°C vs 30°C compared to the other temperatures. Given that *S. pastorianus* CBS 1513 optimal growth temperature is around 25°C (Fischer *et al*. 2016), it is somehow expected that a shift between two temperatures close to its optimal would not induce a drastic difference in the gene expression compared to a shift involving colder temperatures (13°C). Surprisingly, in SD w/o leucine and SD + 6% ethanol, the absolute number of genes differentially expressed is gradually increasing when the shift goes towards the warmest temperature of 30°C. Furthermore, the changes in *S. pastorianus* CBS 1513 transcriptome appeared to be very low in SD media w/o leucine between 13°C and 22°C. It has been shown in a recent study in *Staphylococcus aureus* that amino acid uptake and release is correlated to environmental changes, such as temperature, pH and osmolality, to help the bacteria to adapt (Alreshidi *et al*. 2020). In *S. cerevisiae*, the addition of ethanol induces stress responses and cause cell cycle delay, where heat shock proteins are induced and trehalose is accumulated (Stanley *et al*. 2010). So, when the temperature rises in presence of ethanol, the transcriptional changes may be exacerbated.

**Figure 2:**
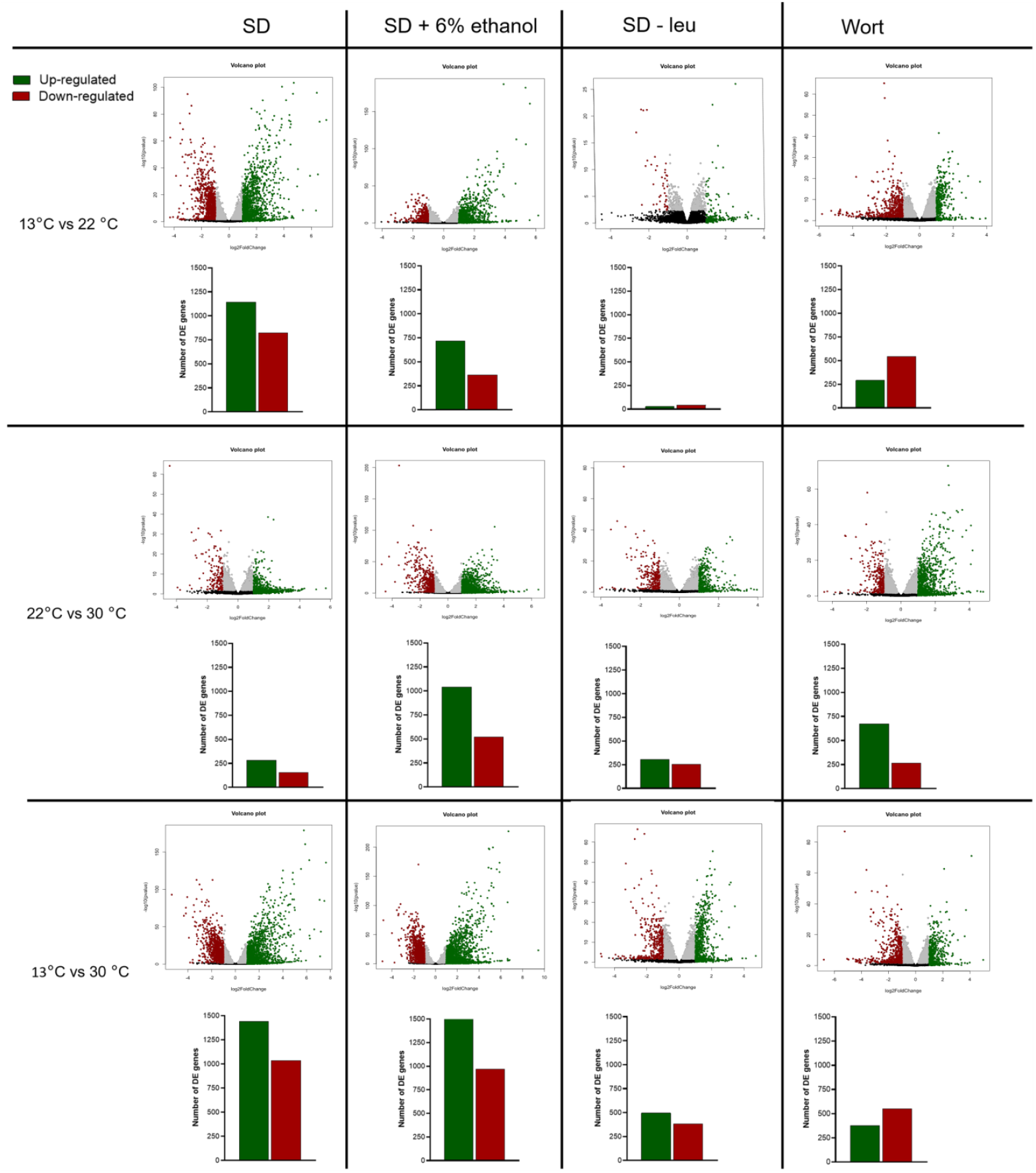
Volcano plots and histograms representing the differential expression and absolute number of DE genes of *S. pastorianus* CBS 1513 genes in SD, SD +6% ethanol, SD w/o leucine and wort media, in each temperature shifts (13°C vs 22°C, 22°C vs 30°C and 13°C vs 30°C). Up- and down-regulated genes are represented in green and red, respectively (absolute fold change > 2 and adjusted p-value < 0.05). Genes with significant, but low expression change (absolute fold change < 2 and adjusted p-value < 0.05) are coloured in grey, and the ones not significant in black (adjusted p-value > 0.05).

### 2. Temperature-dependent media-independent genes are primarily located in the cell wall, plasma membrane and mitochondria

Given that media has an impact on temperature acclimatisation, we isolated and analysed the genes that are temperature-dependent and media-independent. When comparing the expression network at 13°C vs 30°C, 94 temperature-dependent genes were common for all media (Figure 3, Panel A). Using the pathway enrichment tool (yeastMine), we found a significant over-representation of the glycerol degradation pathways. The GO term analysis showed a significant enrichment of mono- and di-carboxylic acids metabolic and catabolic processes, glutamine metabolic process, oxidoreductase activity, vitamin binding, protein folding and co-enzyme binding (Figure 3, Panel B). Several of these temperature-dependent proteins were located in the cell wall, plasma membrane and mitochondrial membrane. Within the core genes we identified *GUT2* (glycerol-3-phosphate dehydrogenase), a mitochondrial gene previously described as partially responsible for cryo-tolerance in *Saccharomyces kudriazevii* (Paget *et al*. 2014). Similar analysis was carried out for the other temperature shifts. We observed that 13 and 29 genes are temperature-dependent and media-independent at 13°C vs 22°C (Figure S3, Panel A) and 22°C vs 30°C (Figure S3, Panel C), respectively. The GO enrichment analysis showed that these genes are mainly involved in transport, regulation of fatty acid metabolic process, short chain fatty-acid catabolism, phosphatidylcholine biosynthetic process, fermentation and oxido-reductions. Seven out of the 13 genes at 13°C vs 22°C are associated with mitochondria subcellular localisation (Figure S3, Panel B and D). This reflects recent findings in laboratory hybrids *S. cerevisiae*/*S. uvarum* where the transcriptional changes at different temperatures were correlated to the parent donating the mtDNA (Hewitt *et al*. 2020). The functional annotation of the temperature-dependent, media-independent genes is presented in Table S3.

**Figure 3:**
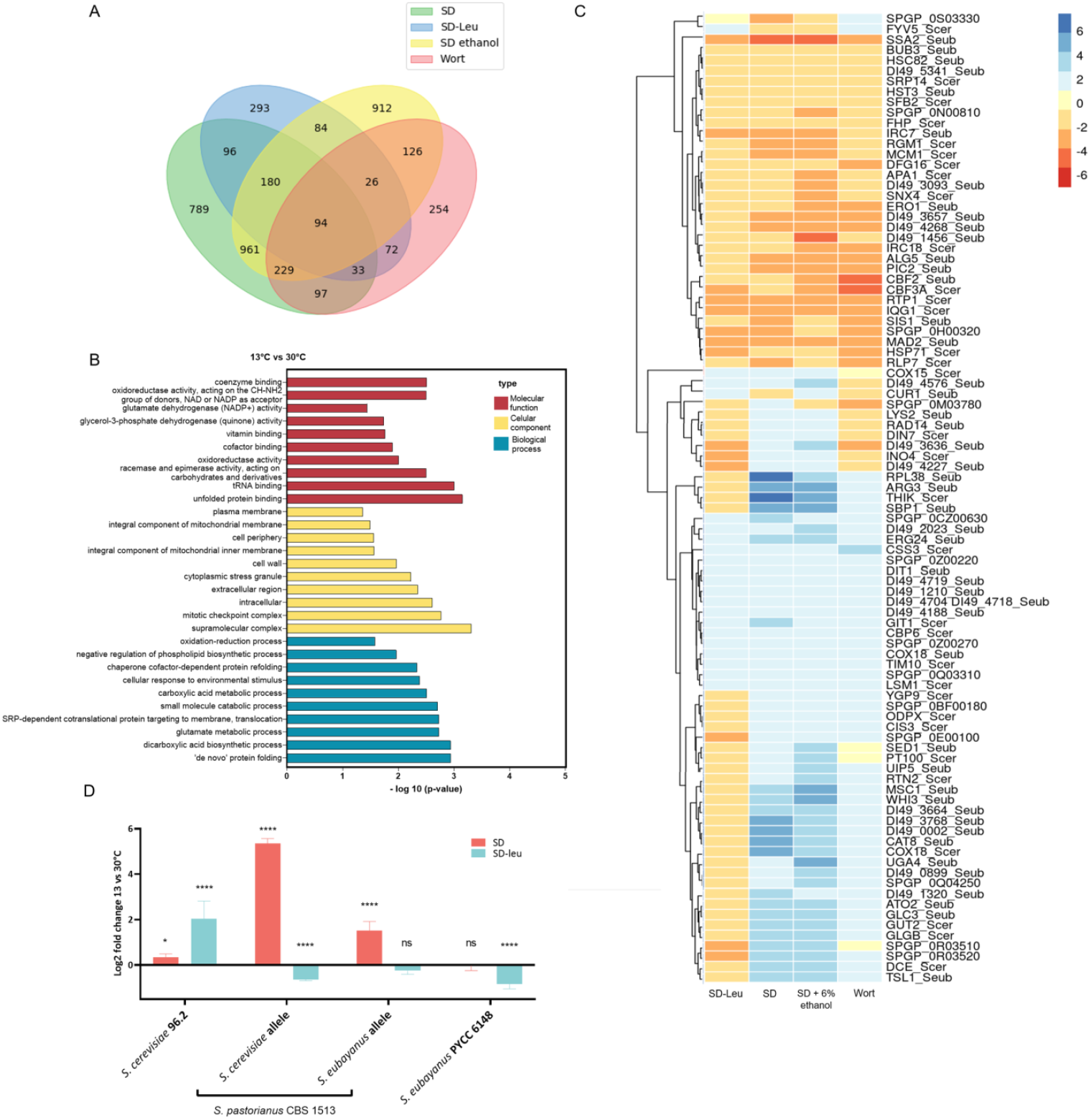
Differentially expressed genes temperature-dependent media-independent. **Panel A**: Venn diagram of the genes differentially expressed between growth at 13°C and 30°C in standard medium (SD; green), standard media without leucine (SD-Leu; blue), standard media with 6% ethanol (SD ethanol; yellow) and maltose rich medium (Wort; pink). The genes present in the intersection of all media conditions are considered temperature-dependent media-independent. **Panel B**: Histogram representing the significance (-log10(p-value)) of the GO terms enriched of the core DE genes at 13°C vs 30°C. Molecular function, cellular component and biological process are coloured in red, yellow and blue, respectively. **Panel C:** Heatmap showing individual expression of the 94 temperature-dependent media-independent core genes at 13°C vs 30°C in SD-Leu, SD, SD+ 6% ethanol and wort. **Panel D:** *GUT2* differential expression at 13°C vs 30°C, for both parental alleles of *S. pastorianus* CBS 1513 (*S. cerevisiae*-like and *S. eubayanus*-like alleles), and for the type strains *S. cerevisiae* 96.2 and *S. eubayanus* PYCC 6148, in SD (red) and SD w/o leucine (blue) media. Error bars denote standard deviations and *p*-values are indicated as: *\* p < 0*.*05 ** p < 0*.*01 ***p < 0*.*001 ****p <0*.*0001;* ns = no significant change.

We identified a small proportion of DE genes in the temperature-dependent group that are affecting growth; such as *DFG16, CIS3* and *WHI3*, involved in invasion during filamentous growth, stability of cell wall, and positive regulation of transcription, respectively. In fact, temperature has an impact on the growth rate through cell division, which makes the biomass yield varying with temperature (Zakhartsev *et al*. 2015). The majority of temperature-dependent genes were non-growth related, such as *CUR1* (involved in cellular response to heat and chaperone binding) and *SSA1* (heat shock protein involved in chaperone cofactor-dependent protein refolding). Indeed, the temperature can trigger the production of chaperones, cold-shock or heat-shock proteins to adapt to the environment (Verghese *et al*. 2012). Temperature can also affect the folding of the proteins, which results in a modified activity. Vitamin biosynthesis, lipid/fatty acid processes and oxido-reduction reactions have also been described in *S. cerevisiae* by Paget and co-workers (2014) to be highly affected by the cold condition. Our transcriptome data in *S. pastorianus* supports this scenario.

In complete SD media, the extracellular leucine uptake is increased at cold temperature as *OPT1* (encoding for an oligopeptide transporter 1 of tetra- and pentapeptides including leucine) is up-regulated. Strikingly, 35% of the 94 core temperature-dependent genes have a unique and opposite behaviour in SD w/o leucine compared to all the other media (Figure 3, Panel C). Such result suggests an impact of leucine uptake on temperature acclimatisation in *S. pastorianus*. Among these genes, *GUT2* is highly up-regulated at 30°C in all media (*i*.*e*. 11, 10 and 4 folds in SD, SD + 6% Ethanol, and Wort, respectively) with the exception of SD media w/o leucine. In this medium, *GUT2* shows an opposite trend as it is 2.6 times less expressed at 30°C. Such differential expression has been validated via RT qPCR for both parental alleles of *S. pastorianus*, and also measured for the parental strains *S. cerevisiae* 96.2 and *S. eubayanus* PYCC 6148 type strains (Figure 3, panel D). *GUT2* expression in medium w/o leucine seems to be hybrid specific as *S. cerevisiae* and *S. eubayanus* parental strains show a different trend: in *S. cerevisiae* 96.2, *GUT2* is up-regulated at 30°C in both media and the *S. eubayanus* PYCC 6148 is not differentially expressed in SD and down-regulated in SD w/o leucine (Figure 3, panel D). Both leucine and *GUT2* are involved in fatty acids regulation, by acting as precursor of branched fatty acids and phospholipids, respectively (Kerkhoven *et al*. 2017, Ferreira *et al*. 2018). Temperature impacts on the physical state of membranes, that needs to stay in a lamellar liquid crystalline phase to function properly. As adaptation mechanism, the yeasts avoid the formation of lamellar gel phase caused by cold temperature by changing the membrane lipids composition, which alters its fluidity (Gunde-Cimerman *et al*. 2014). Short chain lengths and/or unsaturated fatty acids and triacyl-glycerides abundance increase at cold temperatures, while the ratio phosphatidylcholine/phosphatidylethanolamine and the phosphatidic acid content decreases (Martin *et al*. 2007, Redón *et al*. 2011). Here, it is possible that the lack of leucine is already sufficient to limit the amount of phosphatidic acid produced without the need of *GUT2* down-regulation at cold, which would be shifting free glycerol towards glycerone phosphate. In addition, phosphatidylcholine fatty acid is known to be essential for efficient functioning of the mitochondrial Gut2p in *Saccharomyces cerevisiae* (Rijken *et al*. 2007). The absence of extracellular leucine could lead to a decreased production of phosphatidylcholine, which negatively impacts on Gut2p efficiency that is compensated by an upregulation at the transcriptome level at 13°C in SD media w/o leucine compared to 30°C.

We also carried out a similar analysis for the media-dependent, temperature-independent genes (Supplementary File 1). Genes involved in wort utilisation are mainly acting in the super-pathway of glucose fermentation (*ADH1, ADH2, ADH5, GLK1, HXK1, TDH1, PDC5* and *FBP1*). The core temperature-independent genes, triggered by the absence of leucine in the media, are also involved in the super-pathway of glucose fermentation. Interestingly, they are not enriched with leucine biosynthesis biological process, which is concordant with the hypothesis that leucine metabolism is temperature-dependent. Finally, the genes media-dependent temperature-independent involved in ethanol tolerance are enriched with the L-lysine biosynthesis IV pathway and the super-pathway of allantoin degradation, that allows the yeast to use nitrogen as nutrient source by converting the allantoin to ammonia and carbon dioxide.

### 3. Distribution of expression profile in parental alleles shows that *S. eubayanus*-like alleles are primarily involved in temperature response

We broke down the contribution of the specific parental alleles to the expression of the core temperature-dependent DE genes. We first analysed the pool of the 94 temperature-dependent media-independent core genes of which 24 have maintained both parental copies in the genome.

Out of the 53 *S. eubayanus*-like alleles and 33 *S. cerevisiae*-like alleles, there are 40 and 22 alleles that have lost the *S. cerevisiae* and *S. eubayanus* copy, respectively (Table 1). There were also 8 *S. pastorianus*-specific genes (Table 1). Only 3 genes had both alleles DE in the same direction. One DE orthologous pair (*GLC3*) that have both alleles down-regulated at 13°C, encodes for a 1,4-alpha-glucan-branching enzyme a protein involved in the pathway glycogen biosynthesis, previously described to have been affected by temperature in the rainbow trout, human and mouse (Seibert 1985, Naperalsky *et al*. 2010, Hanya and Katz 2018). The two other pairs, *CBF2* and *RTP1*, that encode the centromere DNA-binding part of the protein complex CBF3, and RNA polymerase II assembly factor, respectively, have all their alleles strongly up-regulated at 13°C. These are genes involved in cell division and transcription processes, so it is possible that their change in expression could be due to the different growth rate of the yeast at 13°C and 30°C, rather than the temperature itself. In the core genes involving cold temperature (13°C), there is ∼37% more *S. eubayanus* alleles which are DE, while there is a similar amount of *S. eubayanus*-like and *S. cerevisiae*-like alleles for higher temperatures (22°C vs 30°C) (Table 1). These results suggest that *S. eubayanus* alleles are those who are responding to large temperature shifts and are primarily involved in transcriptional changes at cold temperatures.

**Table 1:** Number of *S. eubayanus-*like, *S. cerevisiae*-like and *S. pastorianus*-specific differentially expressed genes detected in the core genes (temperature-dependent, media-independent) for each temperature condition.

| Condition | *S. eubayanus* -like allele | *S. cerevisiae* -like allele | *S. pastorianus*- specific gene | Total |
| --- | --- | --- | --- | --- |
| 13°C vs 30°C | 53 | 33 | 8 | 94 |
| 13°C vs 22°C | 9 | 3 | 1 | 13 |
| 22°C vs 30°C | 13 | 14 | 2 | 29 |

We expanded this analysis to all DE genes detected in our transcription study to see whether a parental sub-genome is more likely to be differentially expressed under temperature stress. *S. pastorianus* CBS 1513 has evolved from a precursor that appeared after a hybridisation event between a haploid *S. cerevisiae* and a diploid *S. eubayanu*s. There are 5228 and 3751 *S. eubayanus*-like and *S. cerevisiae*-like alleles detected in *S. pastorianus* CBS 1513 genome (Timouma *et al*. 2020). Therefore, if there is no allele bias, out of the total of the DE genes we would expect to identify approximately 58% *S. eubayanus*-like alleles and 42% *S. cerevisiae*-like alleles. We compared the expected and observed number of *S. cerevisiae*-like and *S. eubayanus*-like alleles that show a difference in expression (absolute FC > 1.5 and p-adjusted value <0.05) in all the temperature conditions for each growth media (Table 2). We observed a statistical over-representation of the *S. eubayanus*-like alleles in all the temperature conditions, for all the media except wort for the temperature shifts to 30°C (confidence interval of 95%). In the case of wort there is not a statistically significant difference between the two type of alleles, but there is *ca*. 7 % more DE *S. cerevisiae* alleles than expected (rather than less *S. eubayanus*). This suggest that more *S. cerevisiae* alleles undergo transcriptional changes in wort to help sugar fermentation and to cope with stresses present in this complex rich media.

**Table 2:** Chi-square test on the total number of *S. cerevisiae-*like and *S. eubayanus-*like alleles differentially expressed (absolute fold change > 1.5, p-adjusted value < 0.05) per condition.

| Condition | Media | Total | *S. cerevisiae* -like alleles DE (out of the 3751 *S. cerevisiae* -like alleles present in the genome) | *S. eubayanus* -like alleles DE (out of the 5228 *S. eubayanus* -like alleles present in the genome) | P-value | Outcome |
| --- | --- | --- | --- | --- | --- | --- |
| 13°C vs 30°C | SD | 3885 | 1537 | 2348 | 0.0052 | *S. eubayanus* -like alleles over represented |
|  | SD+6%eth | 4090 | 1628 | 2462 | 0.0106 | *S. eubayanus* -like alleles over represented |
|  | SD-leu | 1838 | 715 | 1123 | 0.0125 | *S. eubayanus* -like alleles over represented |
|  | Wort | 2019 | 802 | 1217 | 0.0615 | No significative difference |
| 13°C vs 22°C | SD | 3357 | 1320 | 2037 | 0.0039 | *S. eubayanus* -like alleles over represented |
|  | SD+6%eth | 2260 | 886 | 1374 | 0.0132 | *S. eubayanus* -like alleles over represented |
|  | SD-leu | 245 | 84 | 161 | 0.0175 | *S. eubayanus* -like alleles over represented |
|  | Wort | 1852 | 703 | 1149 | 0.0009 | *S. eubayanus* -like alleles over represented |
| 22°C vs 30°C | SD | 1193 | 464 | 729 | 0.0436 | *S. eubayanus* -like alleles over represented |
|  | SD+6%eth | 3009 | 1189 | 1820 | 0.0119 | *S. eubayanus* -like alleles over represented |
|  | SD-leu | 1483 | 564 | 919 | 0.0035 | *S. eubayanus* -like alleles over represented |
|  | Wort | 2161 | 864 | 1297 | 0.0909 | No significative difference |

### 4. Analysis of the expression of ortholog parental alleles reveals that the majority of the functionally redundant alleles are not concomitantly expressed by the cell

*S. pastorianus* CBS 1513 strain is maintained in cold temperature for lager beer production, a selective pressure that may impact on its evolution. Group I strains are indeed still evolving, as they show high genomic plasticity and an instable chromosome copy number (Gorter de Vries *et al*. 2020). The higher the loss of *S. cerevisiae* sub-genome in *S. pastorianus*, the lower is the amount of functionally redundant genes (Timouma *et al*. 2020). Here, we investigated how cell response to cold temperatures has shaped the functional redundancy found in *S. pastorianus* CBS 1513. When both *S. cerevisiae*-like and *S. eubayanus*-like alleles are DE, they can either be concordant (both up or down regulated), with or without an allele playing as a major player, or discordant (one allele up the other down regulated). Strikingly, in 81% of the cases, only one of the parental alleles is differentially expressed across all the conditions (Table 3). Out of the remaining 19% of the genes where both parental alleles are DE, 82% have a concordant DE expression (both alleles up or down regulated) across all the conditions, with approximately half of them showing a predominant major player. Only in 3% of the cases the orthologs alleles have a discordant expression (Table 3).

**Table 3:** Expression of the orthologous *S. cerevisiae-*like and *S. eubayanus*-like alleles in the genes differentially expressed for each condition. *S. cerevisiae*-like and *S. eubayanus*-like alleles are annotated “S. cer” and “S. eub”, respectively. Up- and down-regulation, are represented by “+” and “-”, respectively. An allele not DE is annotated “NDE”. Concordant DE expression similar on both alleles is annotated with (+/+ or -/-). If an allele is a major player with more than twice the expression compared to the other allele, it is represented with “++” or “--” for up-regulation and down-regulation, respectively. Discordant expression is annotated with (+/- or -/+).

| Conditions | Media | Total number of pair of othologs with at least one DE allele | DE expression of only one parental allele |  |  |  | DE expression for both parental alleles |  |  |  |  |  |  |  |
| --- | --- | --- | --- | --- | --- | --- | --- | --- | --- | --- | --- | --- | --- | --- |
|  |  |  | Up-regulation of one allele |  | Down-regulation of one allele |  | Concordant DE expression Up regulation |  |  | Concordant DE expression Down regulation |  |  | Discordant DE expression |  |
|  |  |  | S. cer + S. eub NDE | S. cer NDE S. eub + | S. cer - S. eub NDE | S. cer NDE S. eub - | S. cer ++ S. eub + | S. cer + S. eub ++ | S. cer + S. eub + | S. cer - - S. eub - | S. cer - S. eub - - | S. cer - S. eub - | S. cer + S. eub - | S. cer - S. eub + |
| 13°C vs 30°C | SD | 721 | 131 | 173 | 137 | 100 | 17 | 25 | 32 | 13 | 7 | 44 | 23 | 19 |
|  | SD + 6% ethanol | 740 | 147 | 201 | 116 | 103 | 25 | 35 | 39 | 9 | 7 | 29 | 9 | 20 |
|  | SD – Leucine | 279 | 60 | 78 | 51 | 45 | 0 | 4 | 13 | 6 | 2 | 17 | 0 | 3 |
|  | Wort | 290 | 38 | 49 | 78 | 88 | 2 | 3 | 8 | 5 | 3 | 12 | 3 | 1 |
| 13°C vs 22°C | SD | 575 | 125 | 143 | 108 | 88 | 12 | 15 | 28 | 9 | 5 | 18 | 12 | 12 |
|  | SD + 6% ethanol | 316 | 78 | 83 | 56 | 51 | 6 | 14 | 15 | 0 | 0 | 3 | 4 | 6 |
|  | SD – Leucine | 21 | 3 | 2 | 11 | 4 | 0 | 0 | 0 | 0 | 0 | 1 | 0 | 0 |
|  | Wort | 264 | 35 | 39 | 74 | 79 | 1 | 0 | 9 | 4 | 5 | 15 | 2 | 1 |
| 22°C vs 30°C | SD | 148 | 40 | 45 | 26 | 23 | 2 | 1 | 2 | 0 | 1 | 6 | 0 | 2 |
|  | SD + 6% ethanol | 456 | 88 | 147 | 68 | 60 | 12 | 8 | 35 | 7 | 4 | 11 | 8 | 8 |
|  | SD – Leucine | 179 | 43 | 47 | 30 | 40 | 2 | 1 | 3 | 4 | 0 | 8 | 0 | 1 |
|  | Wort | 281 | 69 | 86 | 43 | 34 | 4 | 12 | 17 | 2 | 2 | 4 | 3 | 5 |

The fact that the hybrid can modulate the expression of both types of alleles confers an advantage over the parental strains to achieve the best fitness in a given environment. However, functional redundancy can be costly (and eventually evolutionary unstable) and most of the redundant unused alleles are expected to be silenced or lost as the hybrid is adapting to its new niche. Recent transcription studies on laboratory hybrids *S. cerevisiae* x *S. uvarum* revealed a clear pattern of concerted allelic transcription between the parental alleles (Paget *et al*. 2014). Moreover, a temperature biased gene retention was observed with major loss at 15°C of alleles deriving from the mesophilic parent (Smukowski *et al*. 2019).

### 5. Analysis of the expression of orthologous members of protein complexes shows a trend which favour the formation of either exclusively uni-specific or chimeric complexes

The different nature of protein assemblies can act as an evolutionary force in the hybrid organisms where orthologous members can bind together forming uni-specific or chimeric protein complexes. Considering the presence/absence of parental alleles in *S. pastorianus*, the protein complexes can be either exclusively uni-specific with subunits coming only from one parent (and when all the subunits from the other parent are lost); or exclusively chimeric, with a mixture of subunits from both parents; or partially or fully redundant when a series of protein complexes with different orthologous members can be established because both alleles are present for some or all subunits, respectively. Moreover, according to which alleles are expressed, different scenarios of protein assemblies can occur even when both parental alleles are presents in the genome, specifically in the partially or fully redundant cases (Figure 4, Panel A).

**Figure 4.**
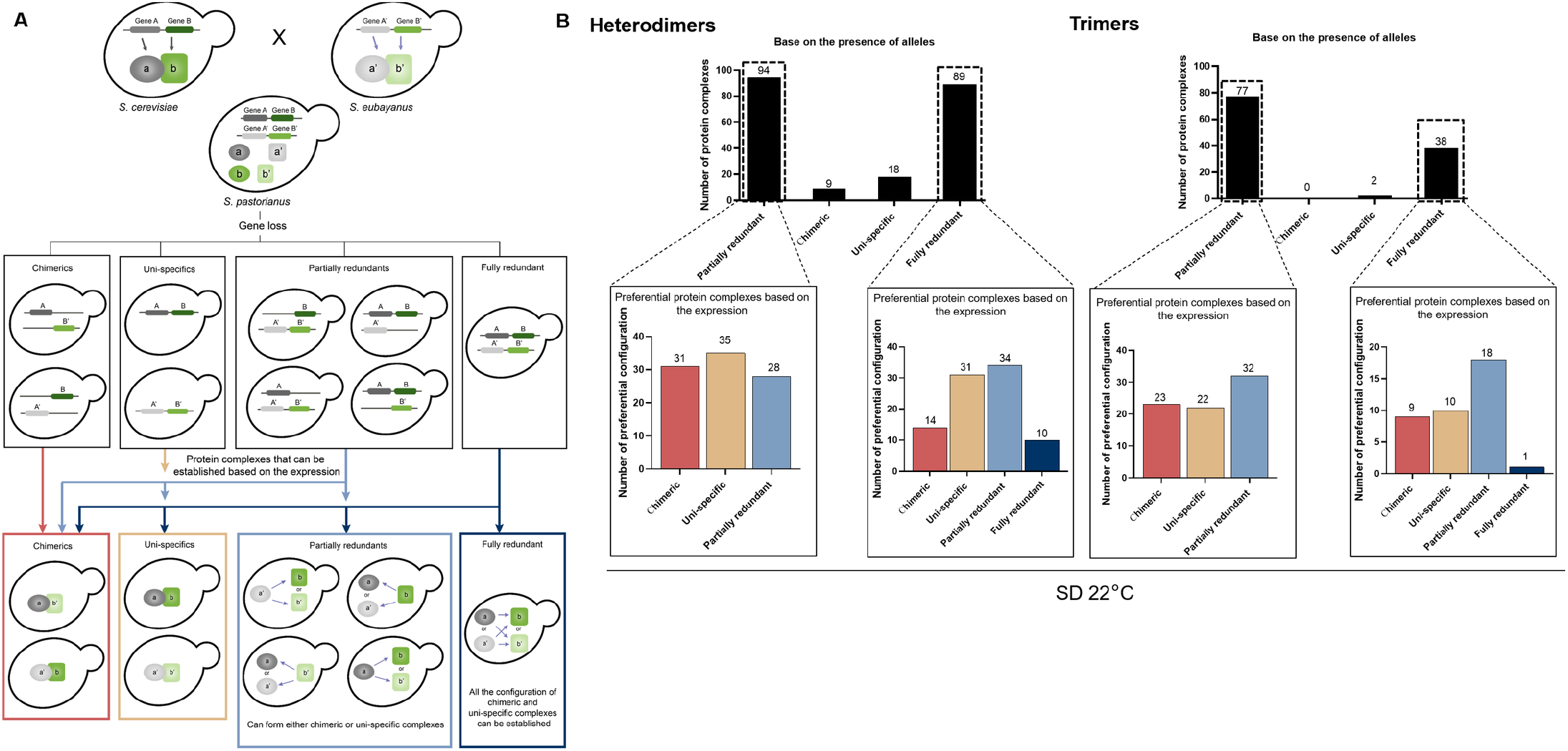
**Panel A:** Protein complexes assembly diagram according to the presence/absence of parental alleles in the hybrid genome. Here, an example of heterodimer is illustrated. After hybridization event, genes A (grey) and B (green) inherited from *S. cerevisiae* and A’ (light grey) and B’ (light green) from *S. eubayanus* are present in the hybrid genome, which evolves through reduction of genetic content (gene loss). When 2 parental alleles are lost, the protein complexes can be either chimeric or uni-specific. When one parental is lost, it creates partially redundant configurations. When all alleles are conserved, it is a fully redundant case. Based on the alleles expression, chimeric and uni-specific can only form chimeric (red arrow) and uni-specific (beige arrow) protein complexes, respectively. Partially redundant cases can form chimeric, uni-specific or partially redundant protein complexes (light blue arrow). Fully redundant cases can form all the configurations (dark blue arrow). **Panel B:** Histogram representing the total number of chimeric, uni-specific, partially and fully redundant configurations in the 210 heterodimers and 117 trimers based on the presence/absence of parental alleles. The partially and fully redundant cases can form different assemblies based on the expression. An example in SD media at 22°C is illustrated.

Given the complexity of the protein-protein network which increases exponentially with the number of subunits, here we restricted our study to homo- and hetero-dimers and trimers (Figure S4, and Table S4). We determined that 575 over the 607 protein complexes originally identified in *S. cerevisiae* are conserved in *S. pastorianus* CBS 1513, with at least one allele identified for each subunit (Table S5). Among these 575 protein complexes, 8 are homodimers, 210 heterodimers and 117 trimers (Figure S4). For these complexes, the majority of orthologous alleles are present and therefore partially or fully redundant protein complexes can potentially be established (Table S6 and Figure 4, panel B). Using the absolute expression, we analysed for each gene which parental allele is the major player in all the environmental conditions studied. An allele has been considered a major player if the expression was at least two times higher than the homologous allele. Such assessment allowed us to recognize which protein assemblies are more prevalent in *S. pastorianus* CBS 1513 when both parental alleles are present (*i*.*e*. partially or fully redundant cases).

Firstly, we analysed the composition of the protein complexes in our standard condition, SD medium at 22°C. Out the 8 homodimers, 3 are exclusively uni-specific as only one type of parental alleles in present in the genome, and 5 are fully redundant. The absolute allele expression showed that 3 out of 5 preferentially express the same parental allele, suggesting that uni-specific protein complexes are more prevalent in these cases (Table S6). The remaining 2 have alleles which are similarly expressed and can therefore form every possible combinations of the complex. In the 210 heterodimers, based on the presence/absence of parental alleles at the genome level, only 9 and 18 complexes can form exclusively chimeric and uni-specific assemblies, respectively, with the remaining being either fully (89) or partially (94) redundant. The absolute expression data for the 89 fully redundant heterodimers suggest that about 53% form predominantly either uni-specific or chimeric complexes (*i*.*e*. 31 uni-specific complexes and 14 chimeric) (Figure 4 Panel B; Table S6). More strikingly, the analysis of the absolute expression of the 94 partially redundant heterodimers, reveal that 70% of the cases are likely to form exclusively chimeric (31) or uni-specific (35) (Figure 4 Panel B, Table S6). In the 117 trimers, based on the genetic, there are no possibility of exclusive chimeric complexes and only 2 cases where the same parental alleles are uniquely retained for all the members of the complex (uni-specific combination). In fact, in the trimers the majority protein complexes are either fully (38) or partially (77) redundant. Similarly, to the heterodimers, the absolute expression data for the 38 fully redundant trimers suggest that 50% forms either exclusively uni-specific or chimeric complexes (*i*.*e*. 10 uni-specific complexes and 9 chimeric) (Figure 4, Panel B; Table S6). Out of the 77 partially redundant cases, 58.5% are more likely to be chimeric (23) and uni-specific (22), respectively (Figure 4 Panel B, Table S6). The same analysis was extended to the other environmental conditions and showed a consistency in this trend (Figure S5).

Overall, these results show that the majority of the protein complexes established in the hybrid are more likely to be either exclusively chimeric or uni-specific, and that the redundancy is discouraged, which is concordant to our previous conclusions at genome scale on the preferential expression of one allele locus (Table 3). This scenario also fits well with the gene balance hypothesis (GBH), which states that the stoichoimetric imbalances can alter protein complexes function, due to the mode of assembly and kinetics (Birchler and Veitia 2010). The GBH predicts that pairs of chromosomes should be co-gained or co-lost to avoid generating imbalanced protein complexes that causes reduced proliferation rates, often observed in aneuploid cells (Chen *et al*. 2019). This could explain our observation that compared to the fully redundant cases, partially redundant cases are more likely to express only one parental allele per subunit, to form uni-specific or chimeric complexes. The level of transcription of orthologous genes/alleles (*cis*-regulation) and/or mutations in up-stream regulatory factors (*trans* effects) could be a way to buffer the concentration of a given subunit (Comai 2005, Tirosh *et al*. 2009). Moreover, as hybridization events bring together diverged genomes within a same nucleus, it has been hypothesized that in addition to the novel epistatic interactions in the hybrid genome, regulatory interferences between the parental sub-genomes may occur, which could lead to a genomic shock, that induce transcriptional changes (Hovhannisyan et al. 2020).

Additionally, for the heterodimers, we investigated how the protein assemblies may change according to the different conditions. In the case of fully redundant heterodimers, there is ca 33% of the cases where the protein complex is preferentially formed with the same parental proteins across all conditions tested, such as, for example, CPX-1277 (Ino2p-Ino4p transcription activation complex) composed by an *S. eubayanus*-like Ino2p and an *S. cerevisiae*-like Ino4p (Figure 5 Panel A). In the case of the 94 partially redundant heterodimers, the expression data shows that ca. 50% of the complexes prefer the same composition across all conditions (Figure 5 Panel B). Thus, it is possible that in these cases the other unused alleles may soon pseudogenise and eventually be lost from the genome. However, there are cases where the nature of the protein complex is different according to the environment, such as CPX-1661 (Spt4p-Spt5p transcription elongation factor complex). In SD medium, this complex that is likely be uni-specific (*S. eubayanus*-like) at 13°C and chimeric (with Spt4 *S. cerevisiae*-like and Spt5p *S. eubayanus*-like) at 30°C. This suggest that some protein complexes may swap parental subunits to adapt to different environmental conditions. A previous study on hybrids of *S. cerevisiae*/*S. mikatae* and *S. cerevisiae*/*S. uvarum* concluded that protein complexes are able to spontaneously exchange orthologous subunits, and that the different types of assemblies have an impact on the phenotype in specific environments (Piatkowska *et al*. 2014).

**Figure 5:**
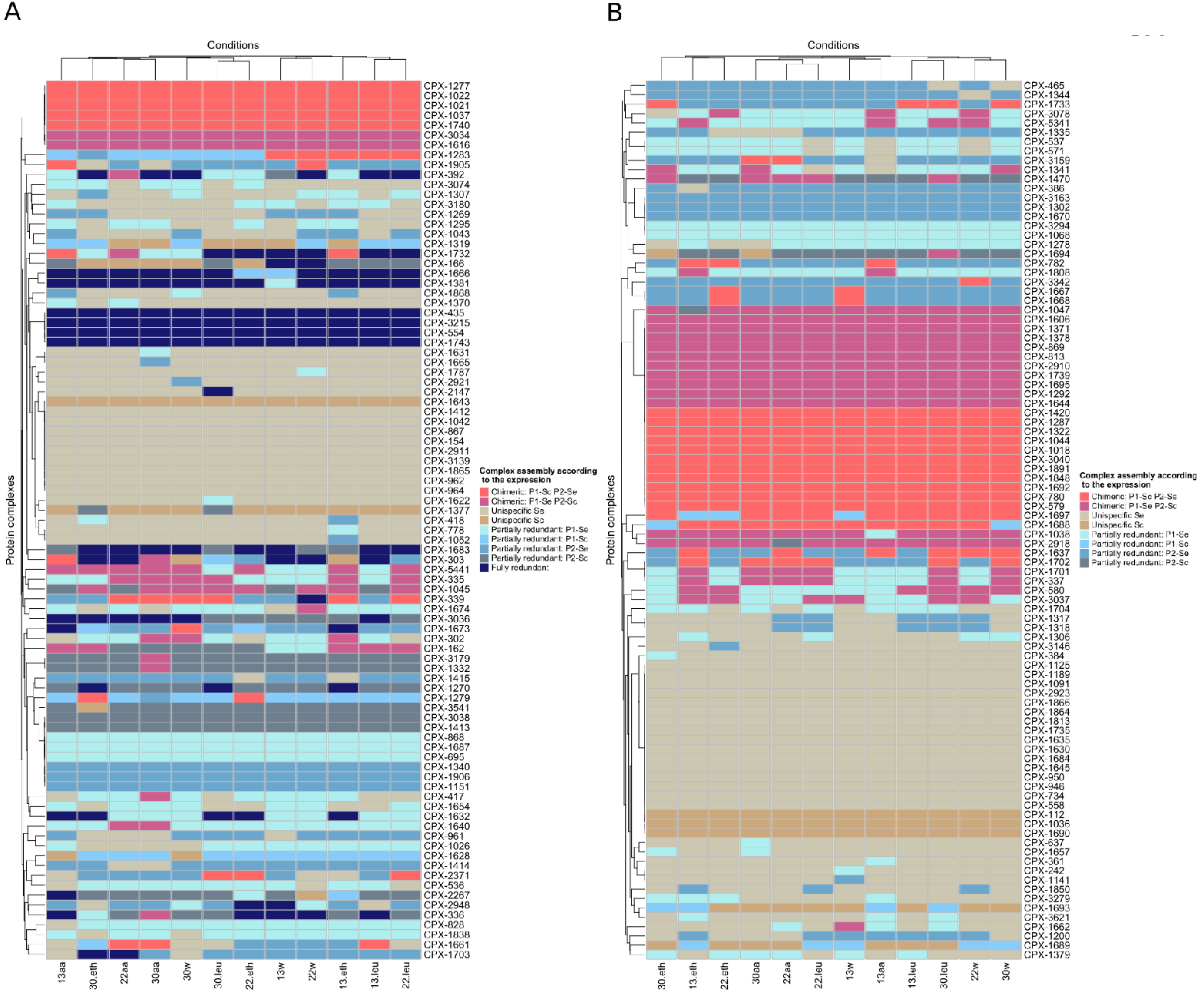
Heatmap showing the assembly of heterodimers (chimeric, uni-specific, partially or fully redundant) based on the absolute expression at 13°C, 22°C and 30°C (prefix “13”, “22”, “30” used in the naming of the conditions, respectively) in SD, SD w/o leucine, SD+ 6% ethanol and wort media (suffix “aa”, “leu”, “eth” and “w” used in the naming of the conditions, respectively). Protein 1 and protein 2 constituting the heterodimer are annotated P1 and P2 in the legend. *S. cerevisiae*-like and *S. eubayanus*-like alleles are abbreviated with “Sc” and “Se” in the legend. **Panel A**: Assemblies of the 89 fully redundant protein complexes. **Panel B**: Assemblies of the 94 partially redundant protein complexes.

## Conclusions

*S. pastorianus*, interspecies hybrid of *Saccharomyces cerevisiae* and *Saccharomyces eubayanus*, is a bottom fermenting yeast that is maintained in cold temperature for lager beer production, a selective pressure that may impact on its evolution. The hybridization event developed unique genetic characteristics for *S. pastorianus*. Here, we analysed transcriptome data from *S. pastorianus* grown at low, ideal, and high temperature and under different media conditions to identify the response of temperature and media acclimatisation at gene expression level. We also identify the protein complex composition and plasticity according to the presence/absence of the parental alleles and the transcriptome profile at different environmental conditions. The primary analysis of the samples clustering suggested that temperature has a higher impact on the transcriptome variability compared to the media. However, the culture media appeared to have an impact on temperature acclimatisation as the absolute number of DE genes at different temperatures varies according to the media. We identified core temperature-dependent media-independent genes. They are mainly involved in protein folding, vitamin biosynthesis, lipid/fatty acid processes and oxido-reduction reactions. We found that extracellular leucine uptake is increased at cold temperature in SD media. Furthermore, the expression of a part of the temperature-dependent genes, including *GUT2*, is affected by the presence/absence of extracellular leucine. We showed that there is an interplay between extracellular leucine uptake and temperature acclimatisation. We also analysed the distribution of expression profiles in parental alleles at the genome level. We strongly predict that overdominance at gene expression level of *S. eubayanus*-like alleles plays a pivotal role for temperature acclimatisation. Expression of ortholog parental alleles reveals that the majority of the functionally redundant alleles are not simultaneously expressed by the cell. We investigated the impact on protein composition when at least one subunit has both parental alleles conserved in the genome. According to our expression data, there is a trend for the formation of either exclusively uni-specific or chimeric complexes. Our results also support the notion that the protein complexes may swap parental subunits to adapt to different environmental conditions.

## Material and Methods

### Genome sequence and annotation

*S. pastorianus* CBS 1513 strain has been sequenced and assembled (Hewitt *et al*. 2014, Okuno*et al*. 2016). Its genome sequence is available from the National Center for Biotechnology Information (NCBI). The Yeast Genome Annotation Pipeline (YGAP) has been used to predict the potential ORFs in its genome (Proux-Wéra *et al*. 2012). HybridMine tool (https://github.com/Sookie-S/HybridMine) has been used to identify the parental allele content in this strain (Timouma *et al*. 2020). Among the 9728 potential ORFs of *S. pastorianus* CBS 1513, HybridMine predicted 5228 *S. eubayanus*-like alleles and 3751 *S. cerevisiae*-like alleles.

### Media and yeast culture

*S. pastorianus* CBS 1513 cells are incubated overnight in Yeast Peptone Dextrose (2% glucose) at 25°C with shaking at 200 rpm. The pre-cultured yeast cells were inoculated in fresh in maltose-rich media (wort), synthetic dextrose (SD) with all amino acids, SD without leucine, SD with 6% ethanol at 13°C, 22°C and 30°C until mid-phase of exponential growth. Liquid fitness assays have been performed for expression validation, three technical replicates for three biological replicates. Cells were grown at 30°C from an optical density of 0.1 (measured at a wavelength of 600nm). Growth was measured every 5 minutes as previously described by Naseeb and Delneri 2012, and recorded by a BMG FLUOstar OPTIMA Microplate Reader (OD 595nm), for up to 55 hours incubation time. Growth parameters were calculated using “Growthcurver” R package.

### Total RNA extraction and quantitative RT-PCR

Total RNA was isolated from three biological replicates of cells collected at mid-log phase (OD_600_ 0.4-0.6) using the RNeasy Mini Kit (QIAGEN, Germany). The lysis was performed by enzymatic digestion of cell wall followed by lysis of spheroplasts. To eliminate genomic DNA contamination, an additional DNAse treatment was performed with RNAse-free DNase set (QIAGEN, Germany) following the manufacturer’s protocol. One microgram of total RNA was reverse transcribed into cDNA using QuantiTect Reverse Transcription Kit (QIAGEN, Germany) according to the manufacturer’s protocol. Optimised qPCR reactions contained 6ng of cDNA, 3pmol each primer and 5 µl of iTAq Universal SYBR Green super Mix 2X in a final volume of 10 µl. Reactions were cycled on a Roche Light Cycler real time System for 35 cycles of: 15 seconds at 95°C; 30 seconds at 55°C; and 30 seconds at 72°C.

### RNA sequencing and data analysis

Total RNA was sequenced using HiSeq4000 Illumina Platform (Genomic Technologies Core Facility, University of Manchester). Quality and integrity of the RNA samples were assessed using a 2200 TapeStation (Agilent Technologies) and then libraries generated using the TruSeq Stranded mRNA assay (Illumina, Inc.) according to the manufacturer’s protocol. A quality control using FastQC was performed on the reads (https://www.bioinformatics.babraham.ac.uk/projects/fastqc/). Trimming and filtering were done using Trimmomatic 0.36 (Bolger *et al*. 2014). Reads were mapped to the annotated genome using STAR aligner 2.5.3a (Dobin *et al*. 2013), and gene expression were counted with the featureCounts tool (Liao *et al*. 2014). Differential gene expression analysis has been done using the DESeq2 package (Love *et al*. 2014) on R version 3.4.4. We applied a false discovery rate (FDR) criterion proposed by Benjamini and Hochberg (Reiner-Benaim 2007) for multiple testing corrections of the raw *P*-value. The threshold of DEGs was set as FDR < 0.05. The genes showing a statistically significant difference in expression between each pair of conditions, and greater than 2-fold change (FC) in expression, were considered as differentially expressed. Data mining has been performed to collect annotation stored in UniProtKB on the *S. pastorianus* CBS 1513 parental alleles identified by HybridMine. A Python 3.6 script that uses the REST application programming interface (API) of UniProt has been developed to query and access its data (Nightingale *et al*. 2017). Gene enrichment has been performed using YeastMine (Balakrishnan *et al*. 2012).

### Protein complexes analysis

Protein complexes present in *S. cerevisiae* has been retrieved from the IntAct Molecular Interaction Database of the EMBL-EBI (Orchard *et al*. 2014). Their presence in *S. pastorianus* CBS 1513 has been detected using an in-house python script (Python version 3.6.9) that searched for each protein in each complex the presence/absence of *S. cerevisiae*-like and *S. eubayanus*-like alleles and their expression in different environmental conditions. When a parental allele is at least two times more expressed than the other, it was considered as major player.

## Aknowledgement

The authors wish to thank Karsten Hokamp and Fiona Mary Roche (Trinity College Dublin) for the useful discussions. This work is supported by the European Commission (grant H2020-MSCA-ITN-2017; number 764364) and BBSRC (BB/L021471/1). LNB is supported by BBSRC (BB/T002123/1) and JP is supported by The National Secretary of Higher Education, Science, Technology and Innovation (SENESCYT, http://siau.senescyt.gob.ec/), Ecuador.

## SUPPLEMENTARY FIGURES

**Figure S1:** Growth curve of *S. pastorianus* CBS 1513 cultured in wort, SD + 6% ethanol, SD media w/o leucine and SD media with all amino acids, at 13°C (Panel A), 22°C (Panel B) and 30°C (Panel C).

**Figure S2:** Validation of DE genes obtained during RNA-seq by RT-qPCRs. Relative mRNA of **Panel A:** *REB1*, **Panel B**: *PRE7*, **Panel C**: *YBR241C*, **Panel D**: *TDA10*, **Panel E**: *POT1* and **Panel F**: *GUT2* at 13°C vs 30°C in SD medium, for both *S. cerevisiae*-like and *S. eubayanus*-like alleles of *S. pastorianus* CBS 1513. Error bars denote standard deviations and *p*-values are indicated as: *\* p < 0*.*05 ** p < 0*.*01 ***p < 0*.*001;* and ns = no significant change upon *t-test*.

**Figure S3: Panel A**: Venn diagram of the genes differentially expressed between growth at 13°C and 22°C in standard medium (SD; green), standard media without leucine (SD-Leu; blue), standard media with 6% ethanol (SD ethanol; yellow) and maltose rich medium (Wort; pink). The genes present in the intersection of all media conditions are considered temperature-dependent media-independent. **Panel B**: Histogram representing the significance (-log10(p-value)) of the GO terms enriched of the core DE genes at 13°C vs 22°C. Molecular function, cellular component and biological process are coloured in red, yellow and blue, respectively. **Panel C**: Venn diagram of the genes differentially expressed between growth at 22°C and 30°C in standard medium (SD; green), standard media without leucine (SD-Leu; blue), standard media with 6% ethanol (SD ethanol; yellow) and maltose rich medium (Wort; pink). The genes present in the intersection of all media conditions are considered temperature-dependent media-independent. **Panel D**: Histogram representing the significance (-log10(p-value)) of the GO terms enriched of the core DE genes at 22°C vs 30°C. Molecular function, cellular component and biological process are coloured in red, yellow and blue, respectively.

**Figure S4:** Histogram representing the total number of protein complex (Y axis) that have X different subunits (X=2* represents homodimers and X=2 heterodimers).

**Figure S5:** Histograms representing the potential assemblies of the partially and fully redundant heterodimers and trimers, based on the expression data obtained at 13°C, 22°C and 30°C, in wort, SD, SD w/o leucine and SD+ 6% ethanol media. Chimeric, uni-specific, partially and fully redundant cases are coloured in red, beige, light blue and dark blue, respectively.

