## Supplementary materials File S1 for "Transcriptional network of the industrial hybrid *Saccharomyces pastorianus* reveals temperature-dependent allele expression bias and preferential orthologous protein assemblies"

**Supplementary File 1:** Media-dependant temperature-independent genes analysis

The media exploited in this study allows us to investigate molecular background behind specific brewing traits of *S. pastorianus* CBS 1513, such as maltose utilisation (wort vs SD media), leucine metabolism (SD w/o leucine vs SD media), and ethanol tolerance (SD + 6% ethanol vs SD media). We identified the genes differentially expressed in these media that are temperature-independant. We identified 303, 202, 167 genes media-dependant temperature-independant involved in nutrients and sugar consumption (wort) (Figure A), leucine metabolism (SD media w/o leucine), and ethanol tolerance (SD media with 6% ethanol).

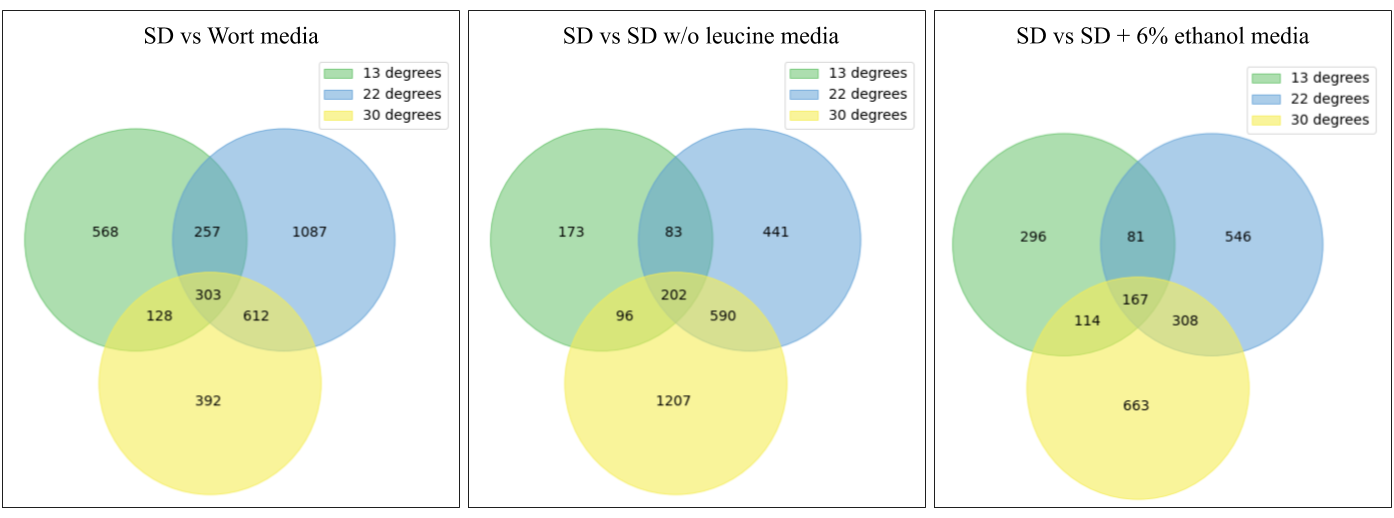
**Figure A:** Venn diagram of the genes differentially expressed between growth in standard medium (SD), standard media without leucine (SD w/o leucine), standard media with 6% ethanol (SD + 6% ethanol) and maltose rich medium (Wort) at 13°C (green), 22°C (blue) and 30°C (yellow).

Genes involved in wort utilisation are mainly acting in the superpathway of glucose fermentation (*ADH1*, *ADH2*, *ADH5*, *GLK1*, *HXK1*, *TDH1*, *PDC5* and *FBP1*). They are associated to small molecule catabolic and biosynthetic process, oxoacid, organic acid, carboxylic acid metabolic and catabolic processes, alcohol biosynthetic process, anion transmembrane transport, cellular amino acid metabolic process. They are mainly engaged in the cytosol, intracellular membrane-bounded organelle and an endoplasmic reticulum. Their molecular function is mainly linked to NAD binding, oxidoreductase activity sugar binding (monosaccharides like glucose), kinase activity (glucokinase, fructokinase, mannokinase), alcohol dehydrogenase [NAD(P)+] activity, and vitamin binding (B6).

The core genes temperature-independant triggered by the absence of leucine in the media are also involved in the superpathway of glucose fermentation (*ADH5*, *GLK1*, *HXK1*, *TPI1*, *ALD5* and *FBP1*). They are linked to gluconeogenesis and hexose (including glucose), oxoacid, carboxylic acid, organic acid and carbohydrates metabolic processes. Interestingly, they are not enriched with leucine biosynthesis biological process, which is concordant with the hypothesis that leucine metabolism is temperature-dependant. These genes are majoritarily located in intracellular anatomical structures, but also cell periphery. They mainly share an oxidoreductase activity, hexokinase activity, symporter activity and secondary active transmembrane transporter activity.

The genes media-dependant temperature independent involved in ethanol tolerance are enriched with the L-lysine biosynthesis IV pathway and the superpathway of allantoin degradation, that allows the yeast to use nitrogen as nutrient source by converting the allantoin to ammonia and carbon dioxide. These genes belong to the small molecule metabolic and catabolic process, carboxylic acid catabolic process, ammonium transmembrane transport and alcohol metabolic process. intracellular anatomical structure and membrane-bounded organelle. saccharopine dehydrogenase activity, catalytic activity, oxidoreductase activity.
