## Supplementary figures and images for "Transcriptional network of the industrial hybrid *Saccharomyces pastorianus* reveals temperature-dependent allele expression bias and preferential orthologous protein assemblies"

### Supplementary Figure S1

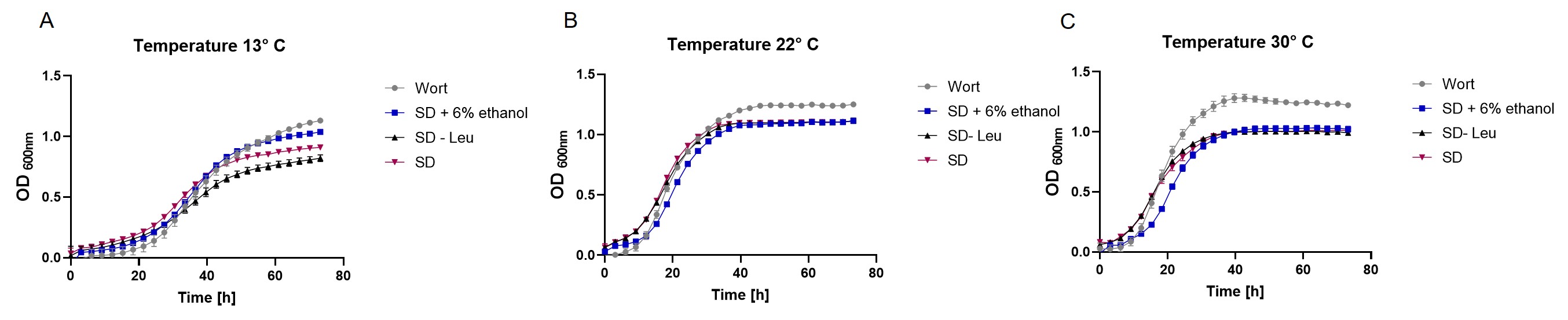

### Supplementary Figure S2

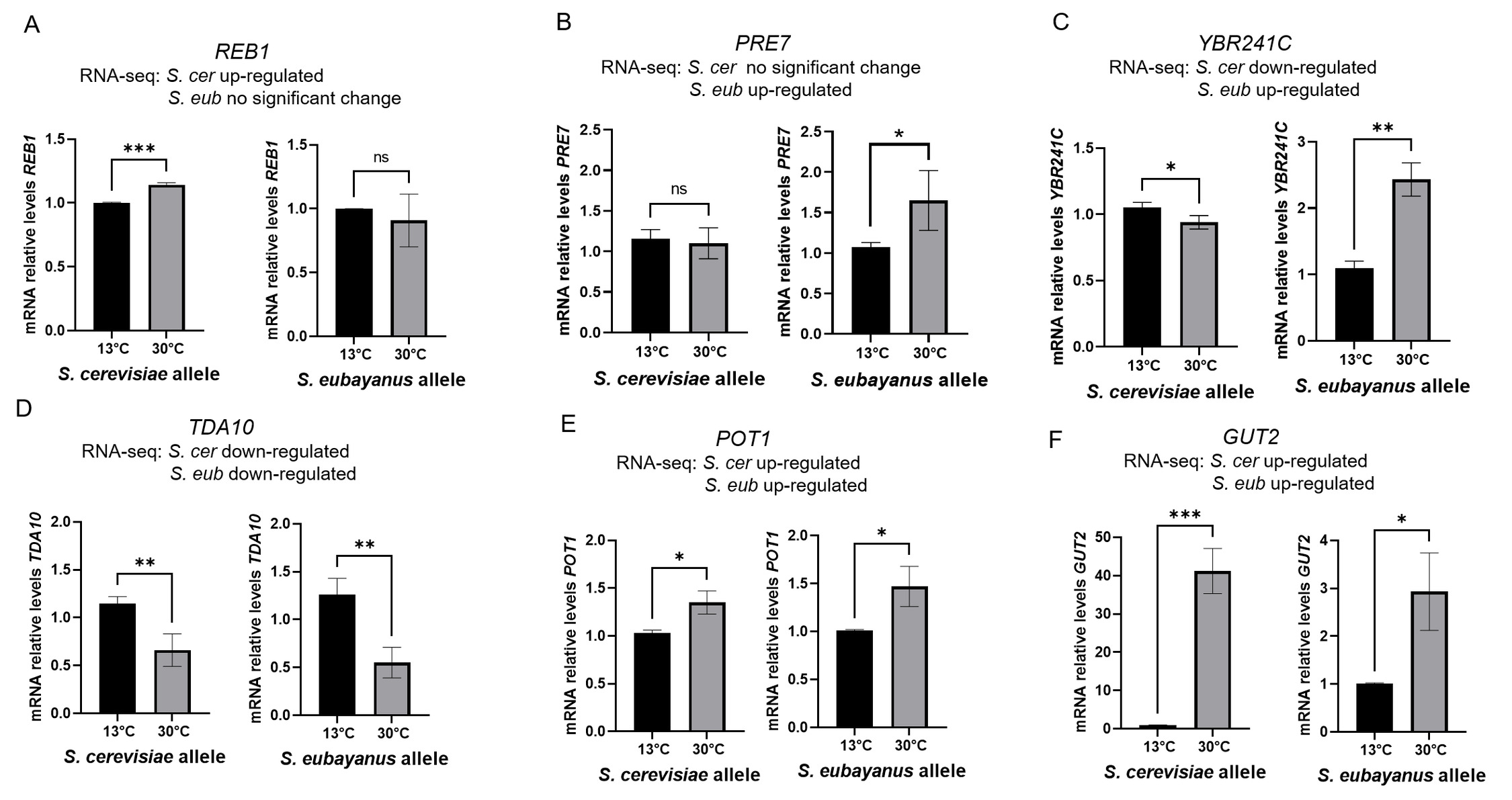

### Supplementary Figure S3

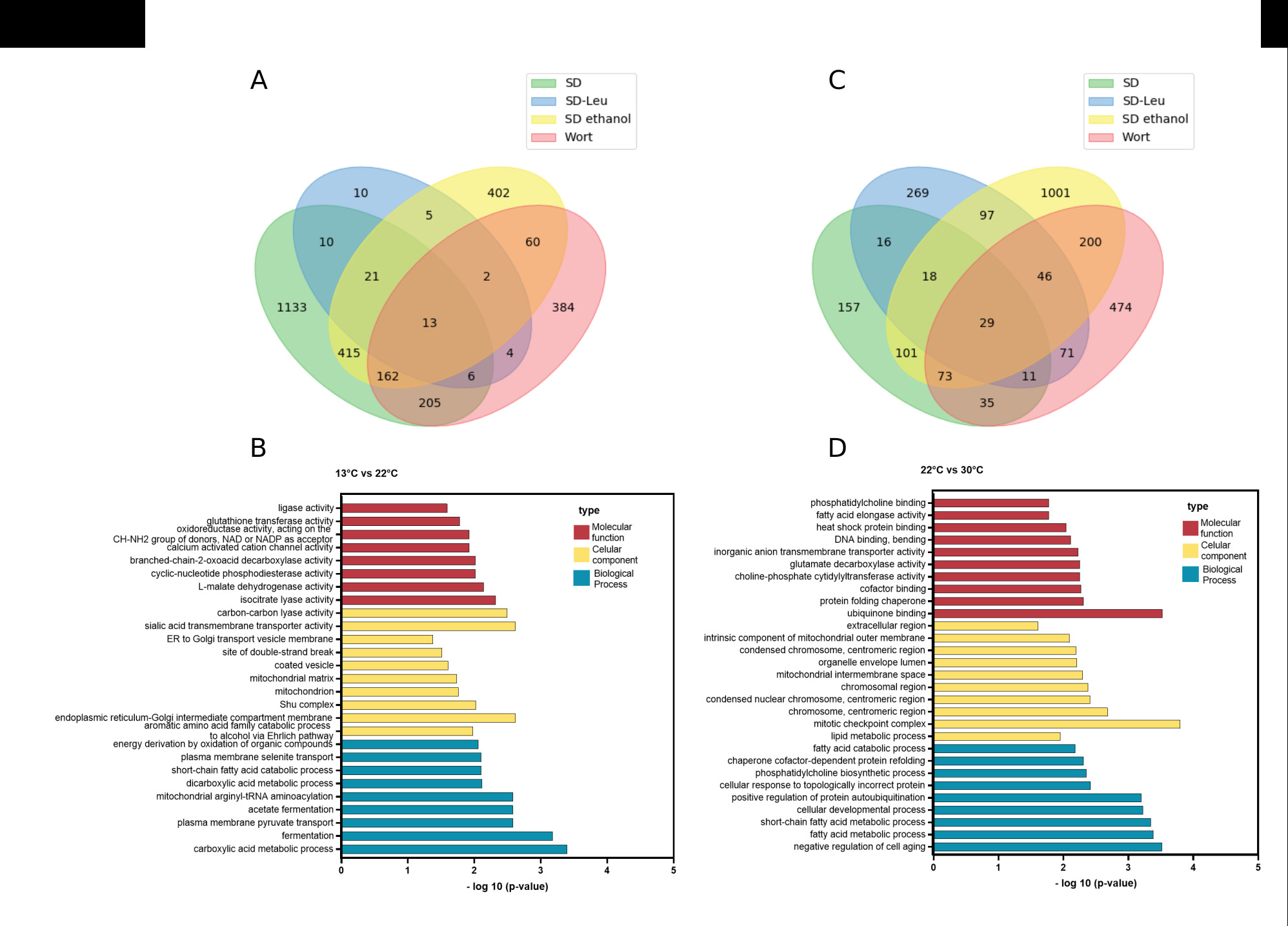

### Supplementary Figure S4

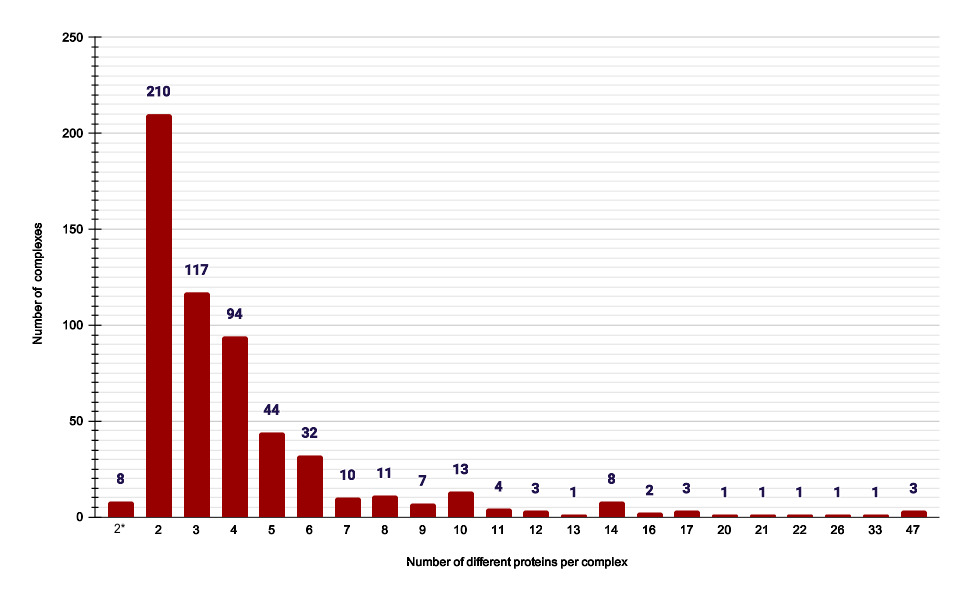

### Supplementary Figure S5

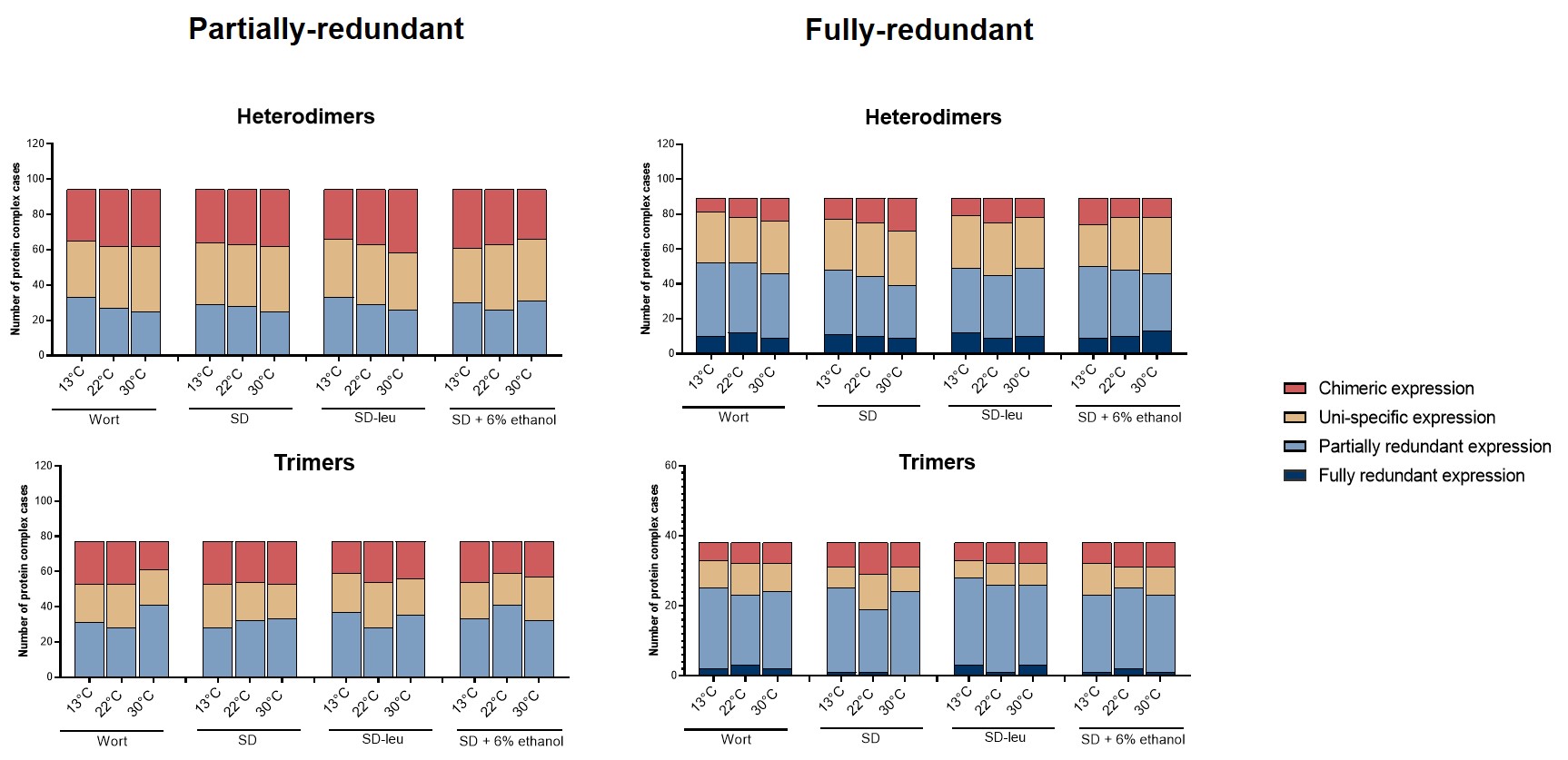
